# Insulin is expressed by enteroendocrine cells during human fetal development

**DOI:** 10.1101/2021.06.07.447234

**Authors:** Adi Egozi, Dhana Llivichuzhca-Loja, Blake McCourt, Lydia Farack, Xiaojing An, Fujing Wang, Kong Chen, Liza Konnikova, Shalev Itzkovitz

## Abstract

Generation of beta cells via transdifferentiation of other cell types is a promising avenue for the treatment of diabetes. Here, we reconstruct a single cell atlas of enteroendocrine cells in the human fetal and neonatal small intestine. We identify a subset of fetal enteroendocrine K/L cells that express high levels of insulin and other beta cell genes. Our findings highlight a potential extra-pancreatic source of beta cells and exposes its molecular blueprint.

---

Identifying insulin-expressing cells outside of the pancreatic islets of Langerhans has important implications for potential cell therapy in diabetes^1^. The pancreas shares its developmental origins with the small intestine, budding from the foregut-midgut boundary at ∼5 weeks gestational age (GA) in humans^2^. Intestinal enteroendocrine cells share many of their transcriptional programs with pancreatic islet cells, including expression of hormones such as somatostatin and ghrelin, as well as common transcription factors. Enteroendocrine cells also share common stimulus-secretion mechanisms with pancreatic beta cells, including glucose-stimulated electrical activities and voltage-gated Ca2+ entry^3^. Several studies demonstrated that perturbations of key transcription factors can result in insulin-expressing enteroendocrine cells. These included the in-vivo inhibition of Foxo1 in Neurog3+ mouse enteroendocrine progenitors^4^ or in human intestinal organoids^5^, as well as the simultaneous induction of Pdx1, MafA and Nuerog3 in the mouse small intestine^6^. Fetal development is often accompanied by plasticity in cell identities^7^. We therefore sought to explore whether insulin might be endogenously expressed in enteroendocrine cells of the human fetus.

We generated a single cell atlas of enteroendocrine cells from human small intestinal samples of four fetal samples (two subjects at 21 weeks GA and two subjects at 23 weeks GA) and two neonatal samples (full term, 2 days old and 3 days old, Figure 1a,b, Supplementary Fig 1a). Enteroendocrine cells clustered into five groups – Enterochromaffin (EC) cells expressing the serotonin catalyzing enzyme Tryptophan hydroxylase 1 (TPH1), Delta cells expressing somatostatin (SST), N/I cells expressing Neurotensin (NTS) and Cholecystokinin (CCK), X cells expressing Ghrelin (GHRL) and K/L cells^3,8^ expressing gastric inhibitory polypeptide (GIP) and the gene encoding glucagon/glucagon-like-peptides (GCG) (Fig 1a-c). We identified distinct markers for each of these cell types (Fig 1c, Supplementary Table 1). We performed differential gene expression analysis, highlighting differences between fetal and neonatal cells (Supplementary Fig 1b-f, Supplementary Table 2).

**Figure 1.**
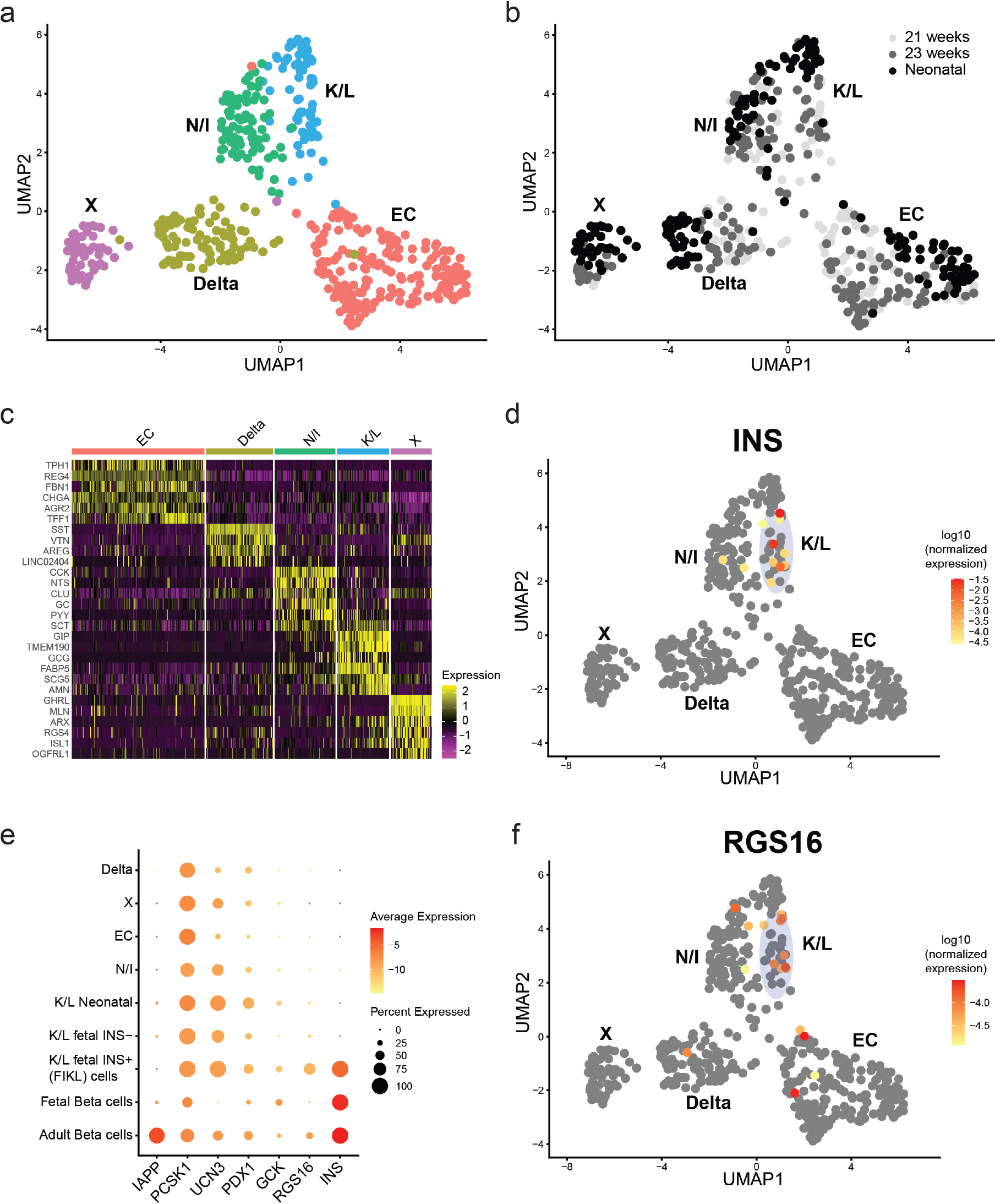
Fetal human K/L enteroendocrine cells contain a subset of INS+ cells. (a) UMAP of human enteroendocrine cells colored by cell type. (b) UMAP of human enteroendocrine cells colored by age. (c) Expression of the top markers for each cell type. (d) INS expression (log10 of the sum-normalized UMI counts). Oval marks the fetal K/L cells, containing 9 out of the 11 INS+ cells. (e) Expression of beta cells genes. K/L cells were split into neonatal, fetal INS-negative and fetal INS+ (FIKL) cells. Fetal and adult beta cell expression signatures were extracted from^9,10^. (f) RGS16 expression (log10 of the sum-normalized UMI counts).

Notably, we identified high expression levels of INS, encoding the insulin protein, in a subset of the fetal K/L cells (9 cells out of 37 fetal K/L cells, hypergeometric test p<4e-11, Fig 1d). We denoted these cells as Fetal INS+ K/L (FIKL) cells. To determine if other beta cell genes were preferentially expressed in FIKL cells, we analyzed the expression of key beta cell genes in FIKL cells and other enteroendocrine lineages as well as in fetal^9^ and adult human beta cells^10^ (Fig 1e, Supplementary table 3). FIKL cells, as well as other enteroendocrine lineages, expressed PCSK1, encoding the proprotein convertase 1, needed for conversion of pro-insulin to insulin. We observed similar broad expression among different enteroendocrine lineages for PDX1, a transcription factor associated with beta cell maturation and insulin transcription, and for UCN3, a marker of beta cell maturation^11,12^. Notably, expression of UCN3 in FIKL cells was higher than in fetal human pancreatic beta cells (Fig 1e, Supplementary table 3). FIKL cells also expressed high levels of GCK, encoding the glucokinase enzyme, that acts as a ‘glucose sensor’ in beta cells and other endocrine cells^13^. In contrast, FIKL cells did not express IAPP, encoding the islet amyloid polypeptide, a hormone promoting satiety that is co-secreted with insulin in adult beta cells^14^ (Fig 1e, Supplementary table 3). Similarly, we observed only low levels of IAPP in fetal pancreatic beta cells (Fig 1e, Supplementary table 3).

The most up-regulated gene in the FIKL cells was RGS16, encoding a regulator of G-protein signaling that has been shown to promote insulin secretion^15^ (Fig 1e, average expression of 4.8e-5 in FIKL cells vs. 5.3e-6 in INS-K/L cells, p<2.5e-5). To further delineate the molecular signature of FIKL cells, we compared the expression of known transcription regulators of adult K or L cell identities^16^ (Supplementary Fig 2, Supplementary Table 2) between fetal and neonatal K/L cells. Among the regulators expressed in our dataset, we found significantly lower expression of both ISL1 (insulin gene enhancer protein, average expression 1.2e-4 in fetal K/L cells vs. 2.7e-4 in neonatal K/L cells, p<0.002) and of PAX6 (paired box-6, average expression 6.1e-5 in fetal K/L cells vs. 1.9e-4 in neonatal K/L cells p<0.0002) and comparable levels of ETV1 (ETS Variant Transcription Factor 1, average expression 1.8e-4 in fetal K/L cells vs. 1.6e-4 in neonatal K/L cells p=0.34).

To validate the existence of INS+ enteroendocrine cells in the fetal intestine, we next used single molecule fluorescence in-situ hybridization (smFISH) of insulin mRNA combined with immunofluorescent antibody staining of the insulin protein^17^ (Fig 2). We hybridized and imaged sections from twelve fetal human subjects of gestational ages between 12W and 23W, as well as sections from three neonatal subjects. We identified multiple cells with high expression levels of insulin mRNAs and proteins in three fetal samples (12W, 14W and 21W) and in none of the neonatal samples. INS+ cells were much scarcer than CHGA+ EC cells (fraction of 0.02+-0.01 of CHGA+ cells, averaged over 41 imaging fields from three fetal samples). INS+ positive cells exhibited a distinct intra-cellular polarization^17,18^, with insulin mRNA localized towards the apical sides of the cells and insulin protein polarized to the basal sides (Fig 2a-b). A similar polarization was previously demonstrated in adult mouse beta cells^17^, and was also apparent when imaging insulin mRNA and protein in adult human pancreatic islet sections (Fig 2c). We further used smFISH to validate co-expression of INS and GCG (Fig 2d, e).

**Figure 2.**
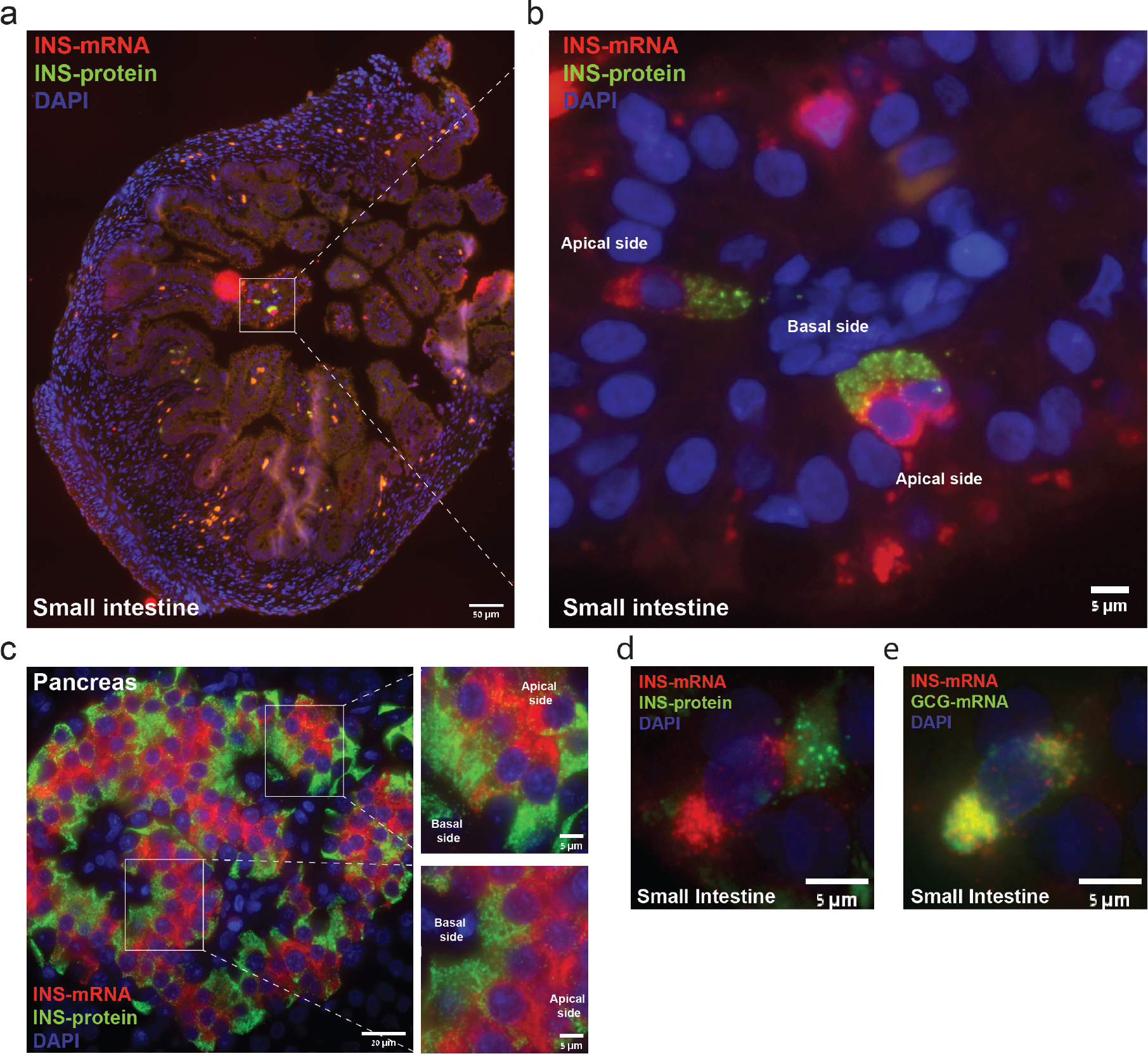
In-situ validation for the expression of INS+ cells in the fetal human small intestine. (a) Intestinal section of 12 weeks GA fetal small intestine simultaneously stained with smFISH probe library for INS-mRNA (red) and an antibody for INS protein (green). Scale bar - 50µm. (b) Blowup of INS+ cells showing the apical polarization of INS-mRNA (red) and basal polarization of INS-protein (green). Scale bar - 5µm. c) Pancreatic human islet (nPOD6447) stained for INS-mRNA (red) and INS-protein (green) demonstrating a similar apical-basal mRNA-protein polarization to that observed in the intestine. Scale bars - 20µm (5µm for blowups). d-e) An INS+ enteroendocrine cell stained with INS-mRNA smFISH probe library (red) and antibody against INS-protein (green, d) and INS-mRNA (red) and GCG-mRNA (green, e). Scale bars - 5µm. In all images nuclei are stained by DAPI (blue).

The physiological role of the intestinal INS+ enteroendocrine cells is unclear, and could either have a local paracrine effect or a systemic endocrine effect. A systemic effect is unlikely, since the numbers of intestinal fetal INS+ cells are around four orders of magnitude smaller than the number of beta cells in the mother’s pancreas (Supplementary note). To assess the potential for local paracrine effects we analyzed the expression of the insulin receptor (INSR) in a recent human fetal cell atlas of the small intestine^19^. We found particularly elevated expression of INSR in enterocytes, enteroendocrine cells and, most prominently in arterial endothelial cells (Supplementary Fig 3). Insulin has been shown to affect vasculature^20^ by stimulating the production of the vasodilator molecule Nitric Oxide (NO) and by promoting vessel sprouting^21^. Insulin can also act as a general growth factor^22,23^, potentially stimulating intestinal tissue expansion at the fetal stages. Importantly, the fact that FIKL cells were not observed in all samples indicates that they might not carry essential physiological functions. Bi-hormonal pancreatic cells, expressing both insulin and glucagon, are commonly seen in the human and mouse fetal pancreas, and are thought to be eliminated, rather than maturing into either alpha cells or beta cells^24,25^. The FIKL cells may be the intestinal equivalent of these pancreatic bi-hormonal cells.

The scarcity of the INS+ enteroendocrine cells currently prohibit secretion measurements in organoids. An open question is what leads to the cessation of insulin expression in the neonatal small intestine. Our analysis uncovered some molecular differences between fetal and neonatal K/L cells, including the robust expression of RGS16 in the fetal INS+ K/L cells and the lower expression of the K/L transcription factors ISL1 and PAX6. The loss of insulin expression post-nataly may be a result of processes occurring at the NEUROG3+ enteroendocrine progenitors, which were not captured in our scRNAseq data. Endocrine cell identity is thought to be shaped by the repression of ‘disallowed genes’, by mechanisms such as microRNAs^26^. It will be interesting to consider potential microRNA regulators that may be differentially expressed in FIKL cells. The induction of insulin-expressing enteroendocrine cell is a promising avenue for cell therapy in diabetes^5^. Our finding of endogenous expression of insulin in fetal K/L cells provides important demonstration of the ability of such cells to surpass lineage barriers towards beta cells and exposes the molecular blueprint that is compatible with a beta cell phenotype.

## Methods

### Intestinal tissue acquisition and storage

Fresh small intestinal (SI) tissue from human neonatal samples were obtained from surgical resections in infants with IRB approval (IRB# PRO17070226). Human fetal SI was obtained through University of Pittsburgh Biospecimen core after IRB approval (IRB# PRO18010491). For single cell sequencing and cryoblocks, tissue was cryopreserved^27^. Briefly, intestinal tissue samples were cut into sub-centimeter pieces and cryopreserved in freezing media (10% dimethyl sulfoxide (DMSO) in fetal bovine serum (FBS)) in a slow cooling container (Mr. Frosty) at -80°C for 24 hours, then transferred into liquid nitrogen for long term storage. For paraffin blocks, tissue was fixed in 4% formalin for 48 hrs, transferred to ethanol until embedded in paraffin. Blocks were stored in paraffin until sectioned for analysis.

### Intestinal Tissue Digestion

Cryopreserved samples were processed per previously published protocol^28^. Briefly, intestinal tissue samples were quickly thawed and washed in RPMI medium plus 10% FBS (Corning), 1X GlutaMax, 10mM HEPES, 1X MEM NEAA, 1 mM sodium pyruvate, 100 I.U/mL penicillin and 100 Pg/ML streptomycin. Next, intestinal tissue was incubated overnight in the same media with 1 Pg/mL DNase 1 and 100 Pg/mL collagenase A. Tissue dissociation was performed on the gentleMACS Octo Dissociator with heaters (Miltenyi Biotec) using the heated human tumor protocol 1. Tissue was then filtered through a 70-Pm nylon mesh cell strainer (Sigma). A single cell suspension was made by washing in DPBS without Ca2+ and Mg2+ (Sigma).

### Single cell RNA sequencing

Single cell suspensions from SI tissue were selected for live cells using a live/dead kit (Miltenyi Biotec). Libraries were made using the Chromium Next GEM Single Cell 3’ kit v3.0 (10X Genomics) with a target of 5,000 cells. 3’ GEM libraries were sequenced at Medgenome. Sequencing was performed on HighSeq lane with two samples per lane^29,30^ or on an S4 NovaSeq lane.

### Single cell RNA sequencing analysis

The single-cell RNAseq data was processed using Cell Ranger 4.0.0 pipeline to align reads and generate count matrix. Cells were background-subtracted as following: background cells were defined as cells with 100-300 UMIs and mitochondrial fraction below 50%. Average expression of background cells was subtracted from all other cells. Subsequently, cells with less than 200 UMIs were filtered out and genes that were expressed in less than 3 cells were removed from data. Single cells data was analyzed using Seurat package^31^ (version 3.2.2). Enteroendocrine single cells were computationally extracted from a larger atlas of fetal and neonatal human small intestinal samples (Egozi et al., in preparation). Enteroendocrine cells were defined as cells with summed expression of established enteroendocrine markers (Supplementary Table 4) above 0.01 of total cellular UMI counts. Cells were further filtered to retain only cells with UMI counts between 10^3^ and 10^3.5^, more than 500 expressed genes and mitochondrial fraction below 30%.

Two clusters that consisted of doublets, containing an enteroendocrine cell and either a T cell or an enterocyte were computationally removed. The cluster of doublets with T cells had a mean expression of T cell markers (Supplementary Table 4) above 0.0035 of total cellular UMI counts, the cluster consisting of doublets with enterocytes had a sum of enterocyte specific markers (Supplementary Table 4) above 0.004 of total cellular UMI counts. Another cluster with upregulated dissociation markers^32^ (above 0.035 of total cellular UMI counts, Supplementary Table 4) was further removed.

For the dot plot in Figure 1e, additional datasets from Cao, J. *et al*.^9^ and Baron, M. *et al*.^10^ were analyzed using Seurat^31^. Cluster annotations were taken from the published metadata and beta cells were retained for the analysis. FIKL cells were defined as fetal K/L cells with insulin expression above zero.

Enrichment of INS+ cells in the fetal K/L cells was calculated using a Hypergeometric test over the fetal cells. Differential gene expression p-values were calculated using Wilcoxon rank-sum tests. P-value for RGS16 levels between FIKL cells fetal INS-K/L cells was calculated using Wilcoxon rank-sum test.

In Supplementary Fig 1, genes with less than 1e-4 normalized expression were removed. P-values were calculated using Wilcoxon rank-sum test. Q-values were computed using the Benjamini-Hochberg false discovery rate correction^33^. In Supplementary Fig 2, data was normalized to the sum of UMIS, p-values were calculated using Wilcoxon rank-sum test. Single cell RNAseq of human intestinal cells in Supplementary Fig 3 was obtained from Elmentaite, R. *et al*.^19^ using cells from the small intestine. Clusters annotations were taken from the published metadata and clusters containing less than 50 cells were filtered out. Expression was normalized using Seurat log normalization - feature counts for each cell were divided by the total counts, multiplied by 10^4^, incremented with a count of 1 and natural-log transformed.

### Immunofluorescence (IF) combined with smFISH

smFISH combined with IF was performed on both paraffin embedded tissue sections and frozen sections, with a modified smFISH protocol that was optimized for human intestinal tissues based on a protocol by Farack, L. *et al*.^17^ and Massalha, H. *et al*.^34^. For frozen sections, intestinal tissues were fixed in 4% Formaldehyde (FA, J.T. Baker, JT2106) in PBS for 1-2 hour and subsequently agitated in 30% sucrose, 4% FA in PBS overnight at 4°C. Fixed tissues were embedded in OCT (Scigen, 4586). 5-8µm thick sections of fixed intestinal tissues were sectioned onto poly L-lysine coated coverslips and fixed again in 4% FA in PBS for 15 minutes followed by 70% Ethanol dehydration for 2h in 4°C.

For paraffin sections, paraffin blocks were cut into 3µm sections. Sections were deparaffinized in Limonene (Sigma 183164) twice for 10 minutes, then moved to 1:1 limonene and Ethanol solution for 10 minutes. Tissues were incubated in 100% ethanol for 5 minutes and then 10 minutes followed by 95% ethanol for 10 minutes. Finally the tissues were transferred to 70% Ethanol for 2h in 4°C.

For both frozen and paraffin sections, tissues were incubated for 10 min with proteinase K in 50°C (10 µg/ml Ambion AM2546) and washed twice with 2× SSC (Ambion AM9765). Tissues were incubated in wash buffer (20% Formamide Ambion AM9342, 2× SSC) for 15 minutes (for forzen sections) or 5 hours (for paraffin sections) and mounted with the hybridization buffer (10% Dextran sulfate Sigma D8906, 30% Formamide, 1 mg/ml E.coli tRNA Sigma R1753, 2× SSC, 0.02% BSA Ambion AM2616, 2 mM Vanadyl-ribonucleoside complex NEB S1402S) mixed with 1:3000 dilution of probes and 1:900 dilution of guinea pig anti-Insulin antibody (Dako, A0564). Hybridization mix was incubated with tissues for overnight in a 30°C incubator. SmFISH probe libraries (Supplementary Table 5) were coupled to Cy5, TMR or Alexa594. After the hybridization, tissues were washed with wash buffer, then incubated with secondary antibody donkey anti-guinea pig Cy3 (1:300, Jackson Laboratories 706-165-148) or goat anti-guinea pig FITC (1:300, Abcam ab6904) for 20 minutes in room temperature.

Tissues were moved to GLOX buffer (0.4% Glucose, 1% Tris, and 10% SSC) containing 50 ng/ml DAPI (Sigma, D9542) for 10 min followed by GLOX buffer wash. Probe libraries were designed using the Stellaris FISH Probe Designer Software (Biosearch Technologies, Inc., Petaluma, CA), see Supplementary Table 5.

### Imaging

Imaging was performed on Nikon eclipse Ti2 inverted fluorescence microscopes equipped with 100x and 60x oil-immersion objective and a Photometrics Prime 95B 25MM EMCCD camera. Image stacks were collected with a z spacing of 0.3-0.5µm. Identification of positive cells was done using Fiji^35^.

## Supporting information

Supplementary table 1-Markers for the five enteroendocrine cell clusters.

Supplementary table 2-Differential gene expression between fetal and neonatal cells for each of the five enteroendocrine cell clusters.

Supplementary table 3-Mean expression of FIKL cells, other enteroendocrine cells and pancreatic fetal and adult beta cells.

Supplementary table 4-Markers used for single cell cluster filtration.

Supplementary table 5-Sequences for the smFISH probe libraries used in this study.

## Data Availability

The data generated in this study is available at the Zenodo repository under the following https://doi.org/10.5281/zenodo.4785359.

## Code Availability

All code used in the paper is available upon request.

---

**Supplementary Figure 1.**
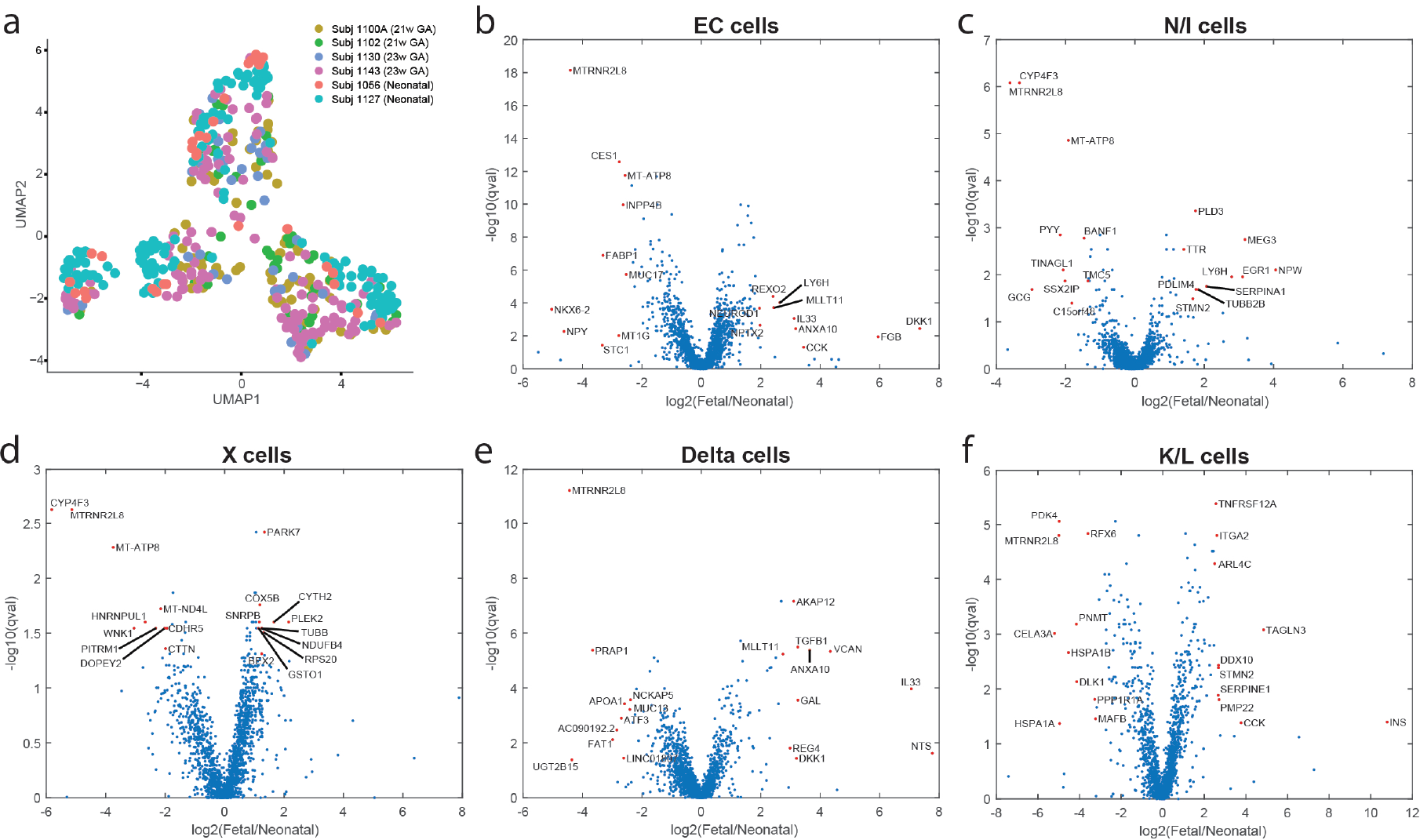
(a) UMAP of the enteroendocrine cell atlas colored by subject. (b-f) Differential gene expression analysis between fetal and neonatal for each cell type. Volcano plots with the 10 genes with highest or lowest expression ratios marked with red dots. X axis is the log2 of the ratio between the average expression in fetal cells and neonatal cells. Y axis is -log10 (q-value), obtained using Benjamini-Hochberg multiple hypotheses correction (Methods).

**Supplementary Figure 2.**
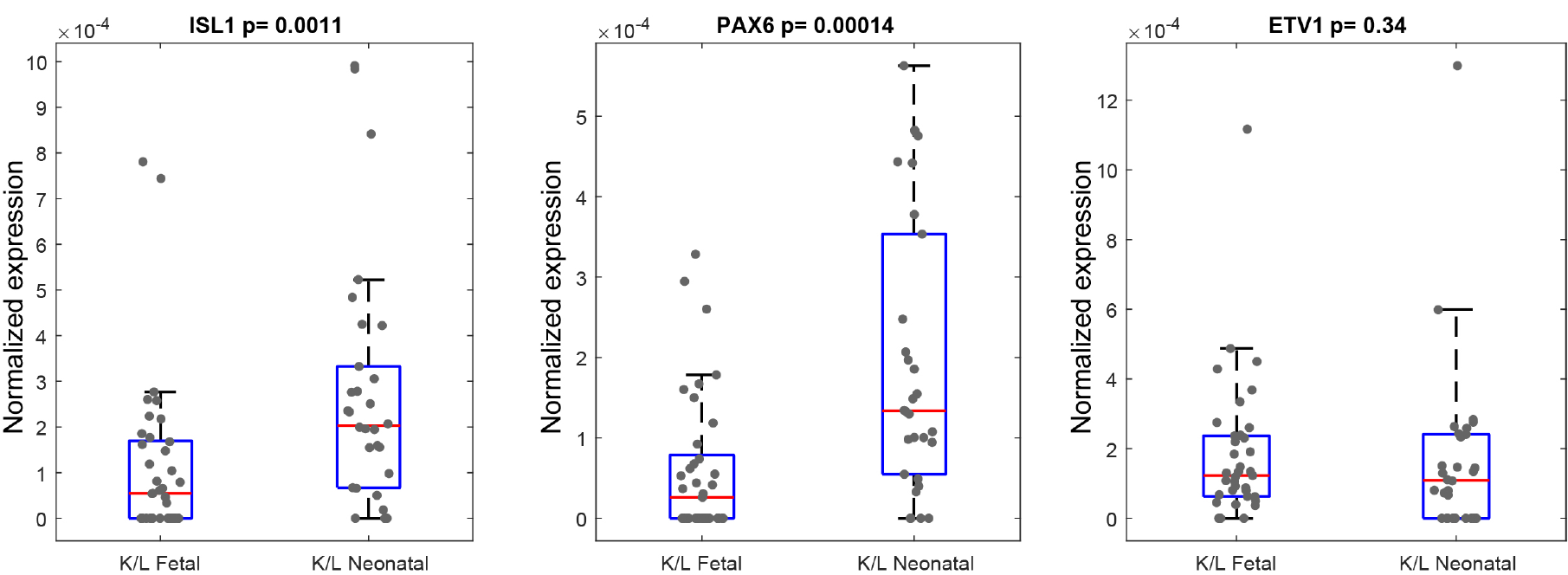
Expression of the transcription factors associated with fate determination of K cells or L cells taken from^16^, included are the only three transcription factors with average expression higher than 1e-5 in either fetal or neonatal K/L cells. P-values calculated using Wilcoxon rank-sum tests. Data normalized to the sum of UMIs.

**Supplementary Figure 3.**
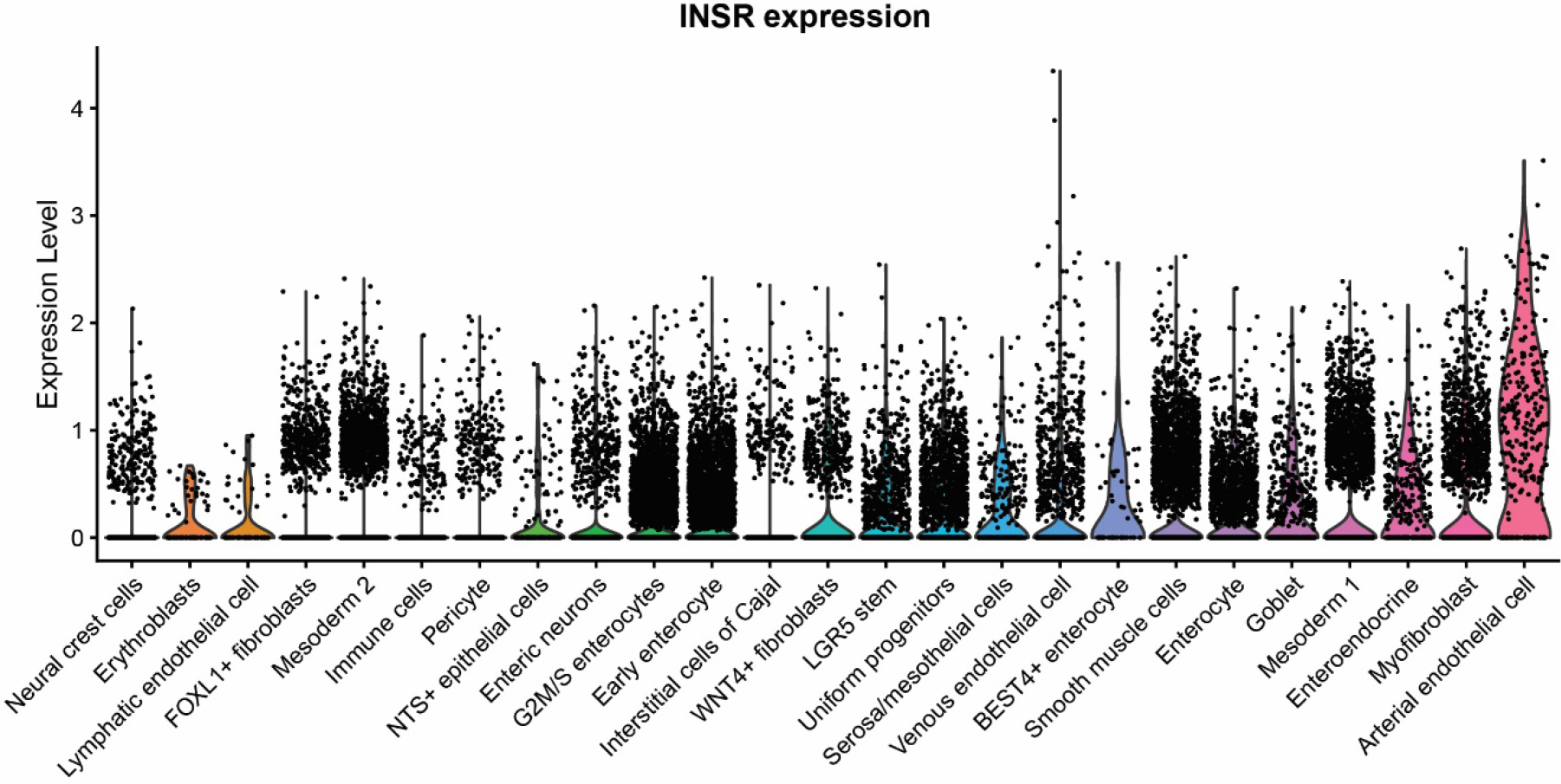
Expression of INSR in small intestinal cell types. INSR expression in all clusters of small intestinal data from^19^. Clusters are ranked by the mean INSR expression. Expression units are Seurat normalized (methods).

## Acknowledgements

We thank Yuval Dor for insightful comments. S.I. is supported by the Wolfson Family Charitable Trust, the Edmond de Rothschild Foundations, the Fannie Sherr Fund, the Dr. Beth Rom-Rymer Stem Cell Research Fund, the Minerva Stiftung grant, the Israel Science Foundation grant No. 1486/16, the Broad Institute-Israel Science Foundation grant No. 2615/18, the European Research Council (ERC) under the European Union’s Horizon 2020 research and innovation program grant No. 768956, the Chan Zuckerberg Initiative grant No. CZF2019-002434, the network of Pancreatic Organ Donors (nPOD) and the Bert L. and N. Kuggie Vallee Foundation and the Howard Hughes Medical Institute (HHMI) international research scholar award. L.K. is supported by previous start-up funds from University of Pittsburgh and current start-up funds from Yale University, Binational Science Foundation award number 2019075, and National Institute of Health R21TR002639 and R21HD102565. No NIH funds were used for the fetal work of these studies.

## Author’s Contribution

S.I, L.K. and A.E conceived the study. D.L. and B.M were involved in sample collection, processing and preparation for single cell analysis. X.A., F.W. and K.C. were involved in library preparation. A.E. performed computational analysis, smFISH+IF experiments and image analysis. L.F. performed smFISH+IF experiment and designed the combined protocol. L.K. and S.I. supervised the entirety of the project. All authors approve of the manuscript.

## Supplementary table captions

**Supplementary table 1 –** Markers for the five enteroendocrine cell clusters, identified by the FindMarkers command in Seurat. Markers included have a log-fold above 0.7 and expressed in at least 0.1 of the cluster cells.

**Supplementary table 2 –** Differential gene expression between fetal and neonatal cells for each of the five enteroendocrine cell clusters. Included are all genes with average expression above 1e-4 of total cellular UMIs in at least one of the two compared populations.

**Supplementary table 3 –** Mean expression of FIKL cells, other enteroendocrine cells and pancreatic fetal^9^ and adult^10^ beta cells.

**Supplementary table 4 –** Markers used for single cell cluster filtration.

**Supplementary table 5 –** Sequences for the smFISH probe libraries used in this study.

## Supplementary note

A typical adult human has 10^9^ pancreatic beta cells^36^. An adult human intestine contains around 10^11^ epithelial cells^37^. Therefore, a fetus at gestation age 21weeks, weighing 420g^38^ would have an order of magnitude of 10^11^*0.420/70= 6*10^8^ intestinal epithelial cells (Assuming the reference man weight of 70kg^37^). In our imaging, we have estimated that the fraction of INS+ cells out of the total intestinal epithelial cells is lower than 1/5000 (around 2% of enteroendocrine cells, which themselves are around 1% of the epithelial cells^39^), yielding an estimated total number of less than 10^5^ INS+ cells throughout the small intestine. This number is four orders of magnitude lower than the numbers of beta cells in the adult pancreas. Intestinal fetal secreted insulin levels would therefore have a negligible effect on the mother’s glucose control, compared to the mother’s pancreatic secretion.

## References

1. Merrell, A. J. & Stanger, B. Z. Adult cell plasticity in vivo: de-differentiation and transdifferentiation are back in style. Nat. Rev. Mol. Cell Biol. 17, 413–425 (2016).

2. Jennings, R. E., Berry, A. A., Strutt, J. P., Gerrard, D. T. & Hanley, N. A. Human pancreas development. Development 142, 3126–3137 (2015).

3. Habib, A. M. et al. Overlap of Endocrine Hormone Expression in the Mouse Intestine Revealed by Transcriptional Profiling and Flow Cytometry. Endocrinology 153, 3054–3065 (2012).

4. Talchai, C., Xuan, S., Kitamura, T., DePinho, R. A. & Accili, D. Generation of functional insulin-producing cells in the gut by Foxo1 ablation. Nat. Genet. 44, 406–412 (2012).

5. Bouchi, R. et al. FOXO1 inhibition yields functional insulin-producing cells in human gut organoid cultures. Nat. Commun. 5, 4242 (2014).

6. Chen, Y.-J. et al. De Novo Formation of Insulin-Producing “Neo-β Cell Islets” from Intestinal Crypts. Cell Rep. 6, 1046–1058 (2014).

7. Suissa, Y. et al. Gastrin: A Distinct Fate of Neurogenin3 Positive Progenitor Cells in the Embryonic Pancreas. PLoS ONE 8, e70397 (2013).

8. Mortensen, K., Christensen, L. L., Holst, J. J. & Orskov, C. GLP-1 and GIP are colocalized in a subset of endocrine cells in the small intestine. Regul. Pept. 114, 189–196 (2003).

9. Cao, J. et al. A human cell atlas of fetal gene expression. Science 370, eaba7721 (2020).

10. Baron, M. et al. A Single-Cell Transcriptomic Map of the Human and Mouse Pancreas Reveals Inter- and Intra-cell Population Structure. Cell Syst. 3, 346-360.e4 (2016).

11. Blum, B. et al. Functional beta-cell maturation is marked by an increased glucose threshold and by expression of urocortin 3. Nat. Biotechnol. 30, 261–264 (2012).

12. Van der Meulen, T. et al. Urocortin3 mediates somatostatin-dependent negative feedback control of insulin secretion. Nat. Med. 21, 769–776 (2015).

13. Matschinsky, F. M. & Wilson, D. F. The Central Role of Glucokinase in Glucose Homeostasis: A Perspective 50 Years After Demonstrating the Presence of the Enzyme in Islets of Langerhans. Front. Physiol. 10, 148 (2019).

14. Hartter, E. et al. Basal and stimulated plasma levels of pancreatic amylin indicate its co-secretion with insulin in humans. Diabetologia 34, 52–54 (1991).

15. Vivot, K. et al. The regulator of G-protein signaling RGS16 promotes insulin secretion and β-cell proliferation in rodent and human islets. Mol. Metab. 5, 988–996 (2016).

16. Gehart, H. et al. Identification of Enteroendocrine Regulators by Real-Time Single-Cell Differentiation Mapping. Cell 176, 1158-1173.e16 (2019).

17. Farack, L. et al. Transcriptional Heterogeneity of Beta Cells in the Intact Pancreas. Dev. Cell 48, 115-125.e4 (2019).

18. Moor, A. E. et al. Global mRNA polarization regulates translation efficiency in the intestinal epithelium. Science 357, 1299–1303 (2017).

19. Elmentaite, R. et al. Single-Cell Sequencing of Developing Human Gut Reveals Transcriptional Links to Childhood Crohn’s Disease. Dev. Cell 55, 771-783.e5 (2020).

20. Muniyappa, R., Montagnani, M., Koh, K. K. & Quon, M. J. Cardiovascular Actions of Insulin. Endocr. Rev. 28, 463–491 (2007).

21. Nowak-Sliwinska, P. et al. Oncofoetal insulin receptor isoform A marks the tumour endothelium; an underestimated pathway during tumour angiogenesis and angiostatic treatment. Br. J. Cancer 120, 218–228 (2019).

22. Susa, J. B. et al. Chronic Hyperinsulinemia in the Fetal Rhesus Monkey: Effects of Physiologic Hyperinsulinemia on Fetal Growth and Composition. Diabetes 33, 656–660 (1984).

23. Menon, R. K. & Sperling, M. A. INSULIN AS A GROWTH FACTOR. Endocrinol. Metab. Clin. North Am. 25, 633–647 (1996).

24. Herrera, P. L. Adult insulin- and glucagon-producing cells differentiate from two independent cell lineages. Development 127, 2317–2322 (2000).

25. Teitelman, G., Alpert, S., Polak, J. M., Martinez, A. & Hanahan, D. Precursor cells of mouse endocrine pancreas coexpress insulin, glucagon and the neuronal proteins tyrosine hydroxylase and neuropeptide Y, but not pancreatic polypeptide. Development 118, 1031–1039 (1993).

26. Kaspi, H., Pasvolsky, R. & Hornstein, E. Could microRNAs contribute to the maintenance of β cell identity? Trends Endocrinol. Metab. 25, 285–292 (2014).

27. Stras, S. F. et al. Maturation of the Human Intestinal Immune System Occurs Early in Fetal Development. Dev. Cell 51, 357-373.e5 (2019).

28. Konnikova, L. et al. High-dimensional immune phenotyping and transcriptional analyses reveal robust recovery of viable human immune and epithelial cells from frozen gastrointestinal tissue. Mucosal Immunol. 11, 1684–1693 (2018).

29. Xin, H. et al. GMM-Demux: sample demultiplexing, multiplet detection, experiment planning, and novel cell-type verification in single cell sequencing. Genome Biol. 21, 188 (2020).

30. Luo, J., Erb, C. A. & Chen, K. Simultaneous Measurement of Surface Proteins and Gene Expression from Single Cells. in T-Cell Receptor Signaling (ed. Liu, C.) vol. 2111 35–46 (Springer US, 2020).

31. Stuart, T. et al. Comprehensive Integration of Single-Cell Data. Cell 177, 1888-1902.e21 (2019).

32. Van den Brink, S. C. et al. Single-cell sequencing reveals dissociation-induced gene expression in tissue subpopulations. Nat. Methods 14, 935–936 (2017).

33. Benjamini, Y. & Hochberg, Y. Controlling the False Discovery Rate: A Practical and Powerful Approach to Multiple Testing. J. R. Stat. Soc. 57, 289–300 (1995).

34. Massalha, H. et al. A single cell atlas of the human liver tumor microenvironment. Mol. Syst. Biol. 16, (2020).

35. Schindelin, J. et al. Fiji: an open-source platform for biological-image analysis. Nat. Methods 9, 676–682 (2012).

36. Scharfmann, R., Staels, W. & Albagli, O. The supply chain of human pancreatic β cell lines. J. Clin. Invest. 129, 3511–3520 (2019).

37. Sender, R. & Milo, R. The distribution of cellular turnover in the human body. Nat. Med. 27, 45–48 (2021).

38. Salomon, L. J., Bernard, J. P. & Ville, Y. Estimation of fetal weight: reference range at 20–36 weeks’ gestation and comparison with actual birth-weight reference range. Ultrasound Obstet. Gynecol. 29, 550–555 (2007).

39. Manco, R. et al. Clump sequencing exposes the spatial expression programs of intestinal secretory cells. Nat. Commun. 12, 3074 (2021).

